# MADloy: Robust detection of mosaic loss of chromosome Y from genotype-array-intensity data

**DOI:** 10.1101/764845

**Authors:** Juan R González, Marcos López-Sánchez, Alejandro Cáceres, Pere Puig, Tonu Esko, Luis A Pérez-Jurado

## Abstract

Accurate protocols and methods to robustly detect the mosaic loss of chromosome Y (mLOY) are needed given its reported role in cancer, several age-related disorders and overall male mortality. Intensity SNP-array data have been used to infer mLOY status and to determine its prominent role in male disease. However, discrepancies of reported findings can be due to the uncertainty and variability of the methods used for mLOY detection and to the differences in the tissue-matrix used. We proposed *MADloy*, the first publicly available software tool that incorporates previous methods and includes a new robust approach, allowing efficient calling in large studies and comparisons between methods. The new method implemented in *MADloy* optimizes mLOY calling by correctly modeling the underlying reference population with no-mLOY status and incorporating B-deviation information. We observed improvements in the calling accuracy with respect to previous methods, using experimentally validated samples, and an increment in the statistical power to detect associations with disease and mortality, using simulation studies and real dataset analyses. We applied *MADloy* to detect the increment of mLOY cellularity in blood on 18 individuals after 3 years, and to confirm that its detection in saliva was sub-optimal (41%). We illustrate the use of MADloy to detect the down-regulation genes in the chromosome Y in kidney and bladder tumors with mLOY, and to perform pathway analyses for the detection of mLOY in blood. *MADloy* is a new software tool implemented in R for easy and robust calling of mLOY status in men aimed to facilitate its study in large epidemiological studies.

## Introduction

The most common somatic mutation in humans is the mosaic loss of chromosome Y (mLOY) in men. Recent evidence suggests the important role of mLOY in numerous diseases, being a biological factor that contributes to overall male mortality (1, 2) and, therefore, is likely to play an important role in male-specific treatments of disease. In particular, mLOY in blood cells increases with age, and is associated with smoking and with the risk of several age-related disorders, including hematological and non-hematological cancers, macular degeneration (3), Alzheimer’s disease (4), major cardiovascular events (5) and suicidal behaviors (6). Despite the accumulating evidence, the mechanisms that trigger mLOY and its clinical consequences are still poorly understood. While susceptibility loci and epigenetic marks for mLOY have been identified (7, 8), larger and more accurate association and functional studies are required; in particular, as some inconsistencies have been observed (9, 10). Therefore, current mLOY calling methods and protocols need to be improved and compared to confirm disease risks and mechanisms. Here, we propose *MADloy* a software tool that incorportates previous methods and implements a novel and robust approach. The software allows comparisons across all current analysis strategies and can be readily used on standard data formats such as PennCNV. We used the novel approach to investigate the optimal processing of mLOY calling from SNP array intensity data and the accuracy of detecting mLOY on different tissue-matrices.

A main source of discrepancy for the associations between mLOY and male diseases can be due to the differences in the methods used to call mLOY status from SNP array intensity data (9, 11). The main signal used for mLOY calling is the log-R-Ratio (LRR) of the SNPs in chromosome Y. As a relative measure of the DNA content of a subject at a genomic locus with respect to a group of individuals, men with mLOY are expected to show low SNP-LRR values across the 56-Mb male-specific region of chromosome Y, which excludes the homologous region between chromosomes X-Y, pseudoautosomal (PAR1 and PAR2) and X-Y transposed (XTR), (12). Computing the mean LLR (mLRR-Y) in the region for each individual Fosrberg et al. called mLOY status on those individuals with mLRR-Y lower than the 99% confidence interval of experimentally induced mLRR-Y variation of normal individuals (1). As a threshold dependent method, this approach, here named *mLRR-Y*_*thresh*_, is sensitive to the characterization of subjects with no-mLOY. Assuming that gains of chromosome Y (GOY) are rare, the positive side of the mLRR-Y distribution (centered at the peak) can be used to identify mLOY outliers by reflecting it to the negative mLRR-Y values to define the calling threshold. However, while GOY events are less frequent than LOY events, they are relevant in tumor tissues. The presence of few of them is enough to affect the mLOY threshold. In addition, the method strongly assumes a symmetrical mLRR-Y distribution, which if not true can also affect the position of the threshold. These two uncontrolled effects can introduce misclassification in mLOY calling, reducing the power to find positive associations (13). Alleviation for mLOY misclassification in the association analyses can be considered if mLRR-Y is taken as a quantitative continuous variable (*mLRR-Y*_*quant*_) (11) or as a proxy of mLOY cellularity (*mLRR-Y*_*cellularity*_) (14). However, these approaches are still limited by the fact that the underlying distribution is a clear mixture of individuals with gains, losses or no changes in chromosome Y genetic content, a similar situation found in copy number variation calling (15, 16). Therefore, improvements on the use of mLRR-Y to call mLOY status should include the robust identification of mLOY events as outliers from an mLRR-Y non-symmetric distribution. In addition, current methods have not used the B-allele frequency (BAF) signal in the homologous region between chromosomes X-Y as an independent signal to confirm findings. The deviation of BAF signal in heterozygous probes from its expected value, namely B-deviation, has been robustly used to detect different types of chromosomal events in mosaicism and compute their cellularity levels (Rodriguez-Santiago, 2010). Taking these issues into account, we have therefore implemented a new mLOY detection method in *MADloy* that integrates robust outlier identification of mLRR-Y values with the use of B-deviation signal. We show the improved performance of the method against current approaches using simulations and real data sets. *MADloy* has been implemented as a Bioconductor package for comprehensive mLOY calling along with visualization functions.

Differences of mLOY detection between blood and buccal smear have also been proposed as a possible source of discrepancy between two large studies that estimated the mortality ratios associated to mLOY (9, 11). To address this issue, we studied 18 individuals with positive mLOY status detected with *MADloy* in blood at base-line and assessed the progression of mLOY cellularity in both blood and saliva in a follow-up visit at 3 years, to determine the extent to which the tissue matrices are comparable for mLOY detection. We finally illustrate the application of *MADloy* in the study of the transcriptional correlates of mLOY in blood and in cancer.

## Methods and Materials

### mLOY calling

The reference methods for calling mLOY status using genotype-array intensity are those described in (1) and (8). Forsberg et al.(1) proposed to analyze the log R ratio (LRR) values of SNPs probes in the male-specific region of chromosome Y (mLRR-Y) in the 56-Mb region between PAR1 and PAR2 on chromosome Y (chrY:2,694,521-59,034,049; hg19/GRCh37). An important consideration is the XTR that is shared between X and Y chromosome is also removed from the analysis because XTR can be affected by alterations in chromosome X, then redefining the mLRR-Y region as: chrY 6,611,498-24,510,581; hg19/GRCh37.

Assuming that mLRR values follow a symmetrical distribution across subjects, individuals are LOY-scored based on the threshold that is defined as the lower limit of the 99% confidence interval of the experimentally induced mLRR-Y variation. Forsberg et al. proposed to model the symmetrical distribution by reflecting the positive values of mLRR-Y over its median (method *mLRR-Y*_*thres*_). LRR is computed as the ratio between two intensities and therefore its distribution is likely to be skewed, in addition the presence of large mLRR-Y values representing XYY gains cannot be fully discarded. Wright et al. (8), therefore, proposed to use mLRR-Y as a continuous variable to measure the degree of mLOY given by its cellularity content (method *mLRR-Y*_*quant*_). However, it is clear that mLRR-Y is a multimodal distribution and ignoring this feature reduces the power in association studies, as it has been seen when copy number variant status is estimated using continuous intensities.

As an alternative method, we proposed to use a robust estimation of the threshold that determines gains and losses in chromosome Y as outliers of of the mLRR distribution. This threshold is estimated by

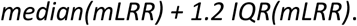

The method is implemented in MADloy together with mLRR-Y_thres_ (1), mLRR-Y_quant_ (8) and mLRR-Y_cellularity_ (14).

To reduce the number false-positive calls, we also propose to combine the information obtained of mLRR-Y with the B-deviation values in X-Y homologous regions which include pseudoautosomal regions PAR1, PAR2, and XTR regions (12) whenever this data is available. The reason to use B-deviation in addition to LRR is because chromosome Y alterations also alter these values in the X-Y homologous regions, due to the allelic imbalance between chromosomes X and Y. We assume an expected BAF of heterozygous probes between 0.45 and 0.55 and by setting a B-deviation threshold of 0.05, we can identify when the BAF is altered. This information can be used as a quality control criterion by identifying those samples classified as LOY or GOY by a mLRR-Y method but with normal values of B-deviation. In addition, samples without alterations in the mLRR-Y values but with alterations in B-deviation classified can provide evidence of technical problems in the processing of the samples. We recommend to visually inspecting discordant samples c and remove cases with clear contamination and other non-LOY events. To this end plots for chromosome X and Y are implemented in the MADloy.

### LRR, mLRR-Y, BAF and B-deviation

mLOY calling is performed on data obtained in PennCNV format for each individual. The format contains SNP, chromosome, position, LRR, BAF and genotype information. LRR and BAF for Illumina arrays (see EGCUT dataset below) can be obtained using GenomeStudio tool (https://www.illumina.com/techniques/microarrays/array-data-analysis-experimental-design/genomestudio.html). For Affymetrix arrays (TCGA dataset), LRR and BAF values are obtained using Affymetrixc Power Tools with the Birdseed v2 algorithm (http://media.affymetrix.com/support/developer/powertools/changelog/index.html) and following the PennCNV-Affy method described in PennCNV webpage (http://penncnv.openbioinformatics.org/en/latest/user-guide/affy/).

For each individual, we computed the 5% trimmed-mean of LRR in autosomes to avoid regions having copy number alterations. The percentage was changed to 25% in the case of cancer samples, where numerous copy number alterations are expected. We then computed the normalized median of LRR-Y (mLRR-Y) with respect to the trimmed-mean of LRR intensity in the autosomes to remove systematic biases across the array. Then, the B-deviation is computed for each sample by selecting the BAF values in the X-Y homologous region (PAR1, PAR2 and XTR) for non-homozygous probes. These probes were selected by setting an upper and lower BAF threshold of 0.8 and 0.2 for normal and variant homozygous genotypes. B-deviation is then calculated as the trimmed mean of the absolute difference between experimental BAF values and 0.5 (i.e expected value) of the heterozygous probes.

### Quality control

We perform a quality control of the samples involved in the analyses by removing those samples with large LRR variability to avoid additional variability likely due to technical artifacts. We followed the Illumina manufacturer’s recommendation of filtering samples with high variable LRR in autosomes (standard deviation of LRR > 0.28). This criterion was used for EGCUT dataset. For TCGA and NIA samples, that were genotyped using Affymetrix, we consider as the normal variability of LRR as 2 times the LRR standard deviation of autosomes.

### MADloy software

We created *MADloy* (https://github.com/isglobal-brge/MADloy), an R package that automates mLOY detection for association studies implementing the methods previously described (17). The package has been submitted to Bioconductor. The core functions include a pipeline to normalize data, perform quality control and summarize data from PennCNV format containing genome-wide information for LRR values. *MADloy* calls mLOY and gains of chromosome Y (XYY) using current methods. Visualization functions are also implemented to help inspect the LRR and BAF values of each sample. A vignette can be found at https://github.com/isglobal-brge/MADloy/tree/master/vignettes.

### Experimental validation

Ten random samples belonging to EGCUT cohort were validated using two Multiplex Ligation Probe-dependent Amplification (MLPA) panels, P070 (MRC-Holland Amsterdam, The Netherlands) covering all subtelomeric regions including the two pseudoautosomal regions (PAR1 and PAR2). Probes targeting the Y chromosome, and a custom-made panel with probes for *SRY* and several autosomal loci, were used to assess the copy number status of chromosome Y with respect to the control loci (autosomal and X chromosome). The MLPA reactions were performed as previously described (18) with the some modifications for custom probe selection (19). We used the relative peak height (RPH) method recommended by MRC-Holland. Non-mosaic losses and gains were expected at relative peak height between (0.5, 1.5) for the pseudoautosomal regions (normally disomic), and between (0, 2) for the Y-unique regions (normally monosomic).

### Statistical analyses

Association studies of mLOY with age and cancer performed in EGCUT and TCGA data were performed using generalized linear models (Gaussian and binomial links, respectively). Models with cancer in EGCUT data were adjusted by age. Transcriptome data belonging to EGCTU data were obtained using HTA 2.0 microarray. Transcriptome of TCGA samples were obtained from RNAseq data available at RTCGA package. Both analyses were performed using *limma* package and the *voom* method was used to obtain continuous data from the RNAseq experiments of TCGA (20). Models were adjusted for surrogate variables using *sva* package to control for experimental differences across samples (21). Enrichment analyses were performed with *GOstats* Bioconductor library (22).

### Simulation studies

We performed a number of simulations to determine the power of the association analyses considering mLOY status as continuous (Wright’s method: mLRR-Y_quant_), as categorical (Forsberg’s: mLRR-Y_thres_ and MADloy’s novel methods). We considered two main scenarios to assess the power to detect significant associations with continuous (age) or categorical (case/control) outcomes. Data were simulated using functions from the CNVassoc package which simulates gains and losses from continuous data (e.g. LRR), considered as surrogate variable of mLOY (23). LRR continuous data was generated using the mean and standard deviation of LRR in normal and LOY cases, and the standard deviation of the outcome using the values observed in EGCUT data. We aimed to simulate the real situation of analyzing the correlation of mLOY with case-control status or age. The effect size varied from 1 to 8, representing age changes for one unit change of mLOY. The sample size also varied (N=200, 300, 500, 750, 1000, 1500). For categorical outcomes, the magnitude of the effect was measured as an odds ratio (OR). We simulated different scenarios with varying mean effects (beta=1.5, 2, 2.5, 3, 3.5, 4) and ORs (1.5, 1.75, 2, 2.5, 3, 3.5, 4).

### EGCUT data

SNP data of 682 individuals from three platforms: OmniX Human370CNV and Metabochip arrays were randomly selected from Estonian Gene Expression Cohort (EGCUT, www.biobank.ee). Selected individuals were older than 18 years of age (mean age 51.4+/-18.2 years). EGCUT comprises a large cohort of about 52,000 samples of the Estonian Genome Center Biobank, University of Tartu (24). Data were genotyped using HumanCoreExome array and all the individuals included in the analysis had a genotyping success rate above 95%. Cryptic relatedness was tested with the PLINK v1.07 software. Only one of each detected relative pair (up to second cousins) was randomly chosen for the detection of genetic mosaicism. Sample mix-ups were corrected using MixupMapper (25). All studies were performed in accordance with the ethical standards of the responsible committee on human experimentation, and with proper informed consent from all individuals tested. LRR and BAF were generated using GenomeStudio software.

### TCGA data

We analyzed 346 paired tumor (n=103) and normal (n=121) male samples of Kidney Renal Clear Cell Carcinoma (KIRC) and Bladder Urothelial Carcinoma (BLCA) belonging to The Cancer Genome Atlas (TCGA) project. Raw data obtained from the Affymetrix Genome-Wide Human SNP Array 6.0 chip were processed with Birdseed v2 algorithm. Clinical data were also downloaded to select male samples and for association analyses. RNAseq data was obtained from RTCGA.rnaseq Bioconductor package that provides counts of 20,532 annotated genes these two tumors.

## Results

### Improved detection of mLOY status

We developed *MADloy*, a bioinformatics tool available as a Bioconductor package, which implements accurate mLOY calling of SNP intensity data using mLRR-Y and B-deviation information across subjects, together with other state-of-the-art methods (http://www.github.com/isglobal-brge/MADloy). For each sample, MADloy first estimates the normalized mLRR-Y given by its ratio with the trimmed-mean of mLRR-Y values in the autosomes. The method uses 5% trimmed-mean to avoid regions having copy number alterations. Calling using mLRR-Y information is given by the identification of unexpected negative values of the mLRR-Y distribution, centered by the expected chromosome Y ploidy for a man (−0.45, one copy), that are lower than 1.2 times the distance of interquantile range. This distance contains ∼%95 of the area under a normal distribution. The interquantile range is robust in the present of outliers and therefore is a suitable measure for simultaneous mLOY and GOY calling. The calling is complemented with B-deviation information in the PAR1/PAR2 regions to reduce the number of false positives of mLOY calls and to estimate their cellularity (see Methods). We then examined whether this new calling method improved calling accuracy and the statistical power in association studies.

We first compared *MADloy* and *mLRR-Y*_*thresh*_ calling methods on 682 males from the Estonian Gene Expression Cohort (EGCUT, www.biobank.ee). Thirty-eight individuals (5.6%) were discarded due to large mLRR-Y variability), see Methods. **Figure 1a** shows the ploidy-centered 5% trimmed mLRR-Y values for each subject with their respective calling status using *MADloy*. **Figures 1b and 1c** illustrate the expected pattern of mLRR-Y and BAF of a normal sample, where the mLRR-Y (blue line) is not far from the expected ploidy value that corresponds to one copy (−0.45) (orange line) and the BAF along the PAR1 (vertical blue rectangle) and PAR2 (vertical yellow rectangle) regions is centered at 0.5 (right Y-scale). **Figures 1b and 1c** show the expected pattern of one sample with mLOY, for which mLRR-Y is lower than the ploidy value and a large BAF split is observed. The size of the split is proportional to the cellularity and is measured by B-deviation defined as the absolute difference between BAF values. The expected value of the B-deviation for heterozygous probes is 0.5 when no mLOY is present.

**Figure 1.**
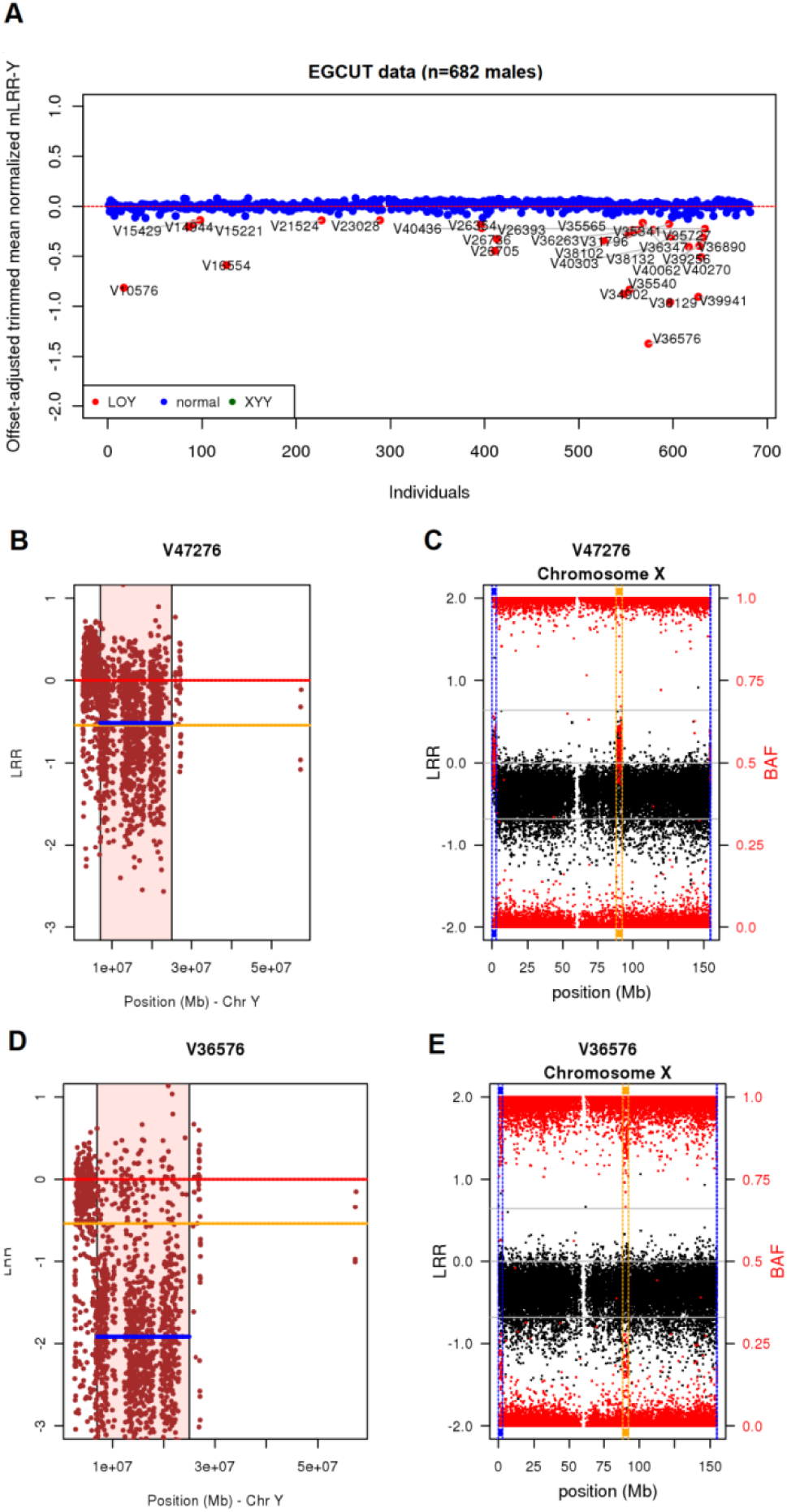
*MADloy* detection in EGCUT data. Figure 1A shows mLOY calling performed using mLRR-Y data by considering our proposed method to robustly detect outliers by considering the possibility of having GOY samples. Figures B and C show the expected behavior of mLRR-Y and BAF data for a normal sample. Figures D and E depicts the expected values of mLRR-Y and BAF data of a mLOY sample. We observe a clear decrease of mLRR-Y (blue line) with respect to the expected value (yellow line) (panel D) and a split BAF in PAR1 (blue box) and XTR (yellow box) regions.

Using *MADloy*, we detected a total of 30 (4.3%) individuals with X-Y allelic imbalance consistent with decreased Y chromosome dosage (**Figure 1a**). Using the *mLRR-Y*_*thresh*_ method, we called 56 samples with mLOY. A main reason for the difference is that the positive distribution of mLRR-Y is less variable than the negative part breaking the symmetry assumption and increasing the negative threshold at which mLOY is called by *mLRR-Y*_*thresh*._ As a result more mLOY individuals are called. **Supplementary Table 1** shows the discordant calls using *MADloy* and *mLRR-Y*_*thresh*_. We investigated the different sources of discordance at the *mLRR-Y*_*thresh*_ calling threshold. For instance, we observed that while V39233 has a lower mLRR-Y value than *mLRR-Y*_*thresh*_, there is no BAF split in PAR1 and PAR2 regions (**Supplementary Figures S1A-B**). In addition, we observed that there are samples with low mLRR-Y due to other causes than mLOY. For instance, **Supplementary Figures S1C-D** shows a case with a BAF split (chromosome X) and LRR values (chromosome Y) consistent with the presence of an additional pair of XX chromosomes, in addition to the XY chromosomes of the individual. These additional X chromosomes are likely due to a contamination of the sample with ∼26% of DNA from another sample.

To increase the experimental support of *MADloy* calling, we validated by MLPA, the positive mLOY status, as detected by *MADloy*, of 10 randomly selected individuals from EGCUT. The results were fully concordant in all ten individuals. **Figure 2A** shows the LRR and BAF of probes across chromosomes Y and X for one individual. **Figure 2B** illustrates the validation of two mLOY cases (V32199 and V40611) with different cellularity (30% and 52%, respectively). We observed that for individuals with mLOY the relative peak heights (RPH) of Y probes were clearly reduced compared to the RPH of the X probe (**Figure 2B**).

**Figure 2.**
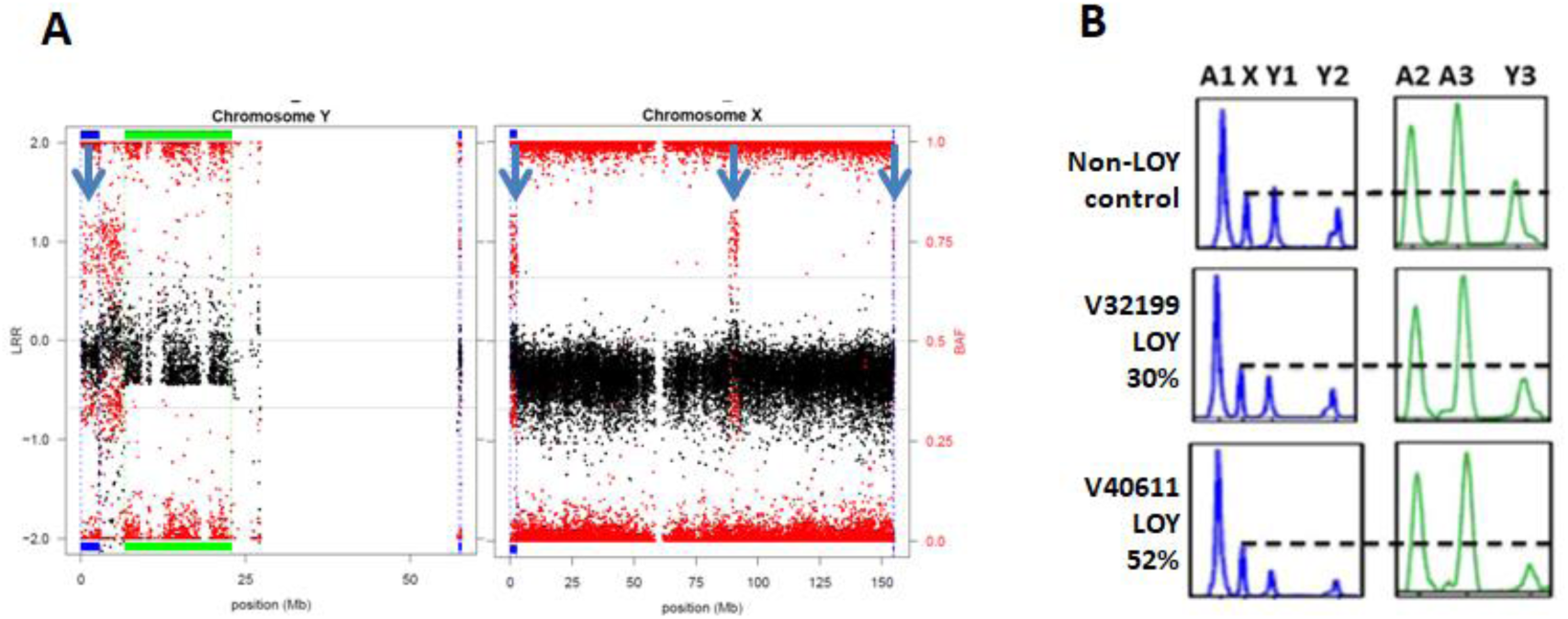
Experimental mLOY validation. **A)** LLR (black) and BAF (red) values of probes along chromsomes Y and X for an individual with positive mLOY status. Low values of LLR-Y are observed except for the PAR1 and PAR2 regions where a clear BAF split is observed. The BAF split is confirmed in the X chromosome. **B)** MLPA validation of two individuals with mLOY with different cellularity content, compared with a normal individual (top). Two MLPA panels are shown for each individual, showing relative peak height (RPH) for probes in autosomes (A1, A2, A3) and in the X and Y chromosomes (Y1, Y2, Y3). Marking the peak for the chromosome X probe (dotted line), the figure shows a clear reduction in the RPH values of Y probes for the individuals with mLOY.

To address the differences in mLOY detection across different tissues, we asked the extent to which mLOY status in blood could be detected in buccal smear. 18 individuals from EGCUT, who were detected with mLOY using *MADloy* in blood, were randomly selected and re-contacted after 3 years to assess the progression of mLOY in blood and evaluate its detection by *MADloy* in buccal smear (**Table 1**). We observed that mLOY in blood persisted in time (**Table 1)**. Detected from BAF signals, the estimated proportion of cells with mLOY increased in 16 cases (88%) while it decreased in 2 (12%). In saliva, we discarded one individual for low quality of the sample and observed mLOY in only 7 individuals (41%). All cases of mLOY in saliva showed lower cellularity than mLOY in blood at followup. Using the same detection methods and procedures, we therefore confirmed that the detection power of mLOY in blood is substantially decreased if assessed in buccal smear.

**Table 1:** Cellularity evolution of LOY detected in blood of 18 individuals after 3 years. At follow-up mosaicism was also tested in saliva (ND: Not Detectable).

|  | Baseline detection |  | Follow-up at 3 years |  |
| --- | --- | --- | --- | --- |
| Sample | Blood DNA | % cells | Blood DNA (% cells) | Saliva DNA (% cells) |
| V15429 | Y loss | 27 | 30 | ND |
| V16554 | Y loss | 52 | 67 | 57 |
| V16763 | Y loss | 41 | 50 | ND |
| V19232 | Y loss | 41 | 53 | ND |
| V22330 | Y loss | 52 | 46 | ND |
| V23235 | Y loss | 37 | 48 | 27 |
| V26736 | Y loss | 42 | 49 | ND |
| V31014 | Y loss | 46 | 48 | 30 |
| V31220 | Y loss | 29 | 39 | 17 |
| V32199 | Y loss | 30 | 40 | ND |
| V32632 | Y loss | 46 | 82 | ND |
| V32752 | Y loss | 32 | 35 | ND |
| V32850 | Y loss | 33 | 43 | ND |
| V34568 | Y loss | 37 | 44 | ND |
| V37288 | Y loss | 31 | 44 | ND |
| V40611 | Y loss | 52 | 29 | 32 |
| V44342 | Y loss | 35 | 37 | 20 |
| V47558 | Y loss | 37 | 39 | 18 |

### mLOY improves the statistical power of association studies

Using a series of simulations, we first compared *MADloy* performance with mLRR-Y_thresh_ and mLRR-Y_quant_. The simulations were performed using an independent simulator for copy number variation, CNVassoc assuming that the distribution of normal and mLOY cases, with the proportions observed for in the EGCUT study, followed those observed for CNVs. We studied the power to detect true associations between mLOY and a quantitative trait. mLOY calling was performed with MADloy and Fosberg’s methods and associations were also assessed with mLRR-Y as a quantitative variable (**Supplementary Figure S2**). We found that *MADloy* had the largest statistical power in all scenarios while mLRR-Y_quant_ was the least powerful. In particular, we observed that an association of magnitude 2 per quantitative trait unit was detected with using *MADloy* with a power of 80% in 1,000 individuals while for *mLRR-Y*_*thresh*_ a magnitude of >2.2 was required for the same power. We also observed similar results for dichotomous traits in case-control studies (**Supplementary Figure S3**). The differences in power were given by a lack of symmetry in the mLRR-Y distribution that resulted in misclassification by *mLRR-Y*_*thresh*_. In addition, the simulations confirmed that treating mLRR-Y as quantitative when the underlying distribution is a mixture is a suboptimal approach.

We then compared in real setting the associations with age and cancer risk with mLOY called with MADloy and *mLRR-Y*_*thresh*_ in EGCUT. In this case we also aimed to evaluate the effect incorporating B-deviation signal in *MADloy* calling. We confirmed the association between mLOY and age. We found that mLOY called by *MADloy* with B-deviation showed the strongest and most significant association with age (beta=21.21, P=2.0×10^−10^). However, when mLOY status was called using mLRR-Y information only, either by *MADloy* or by *mLRR-Y*_*thresh*_, the effect of association was lower (**Figure 3A)**. Note that the largest impact in the association was when considering *mLRR-Y*_*thresh*_ method, suggesting again that the symmetry assumption is less accurate than the interquantile range to define the mLOY calling threshold. We observed analogous results for the association between mLOY and cancer in EGCUT. For MADloy with and without B-deviation we observed similar significant increased risks of any tumor in models adjusted by age (OR=2.89, P= 0.0390, OR=2.86, P= 0.0367). However, for *mLRR-Y*_*thresh*_, we did not find statistically significant differences (OR=1.84, P=0.1720), confirming that the loss of power by methods that do not consider the possibility of having GOY nor the B-deviation can affect statistical significance (**Figure 3B**).

**Figure 3.**
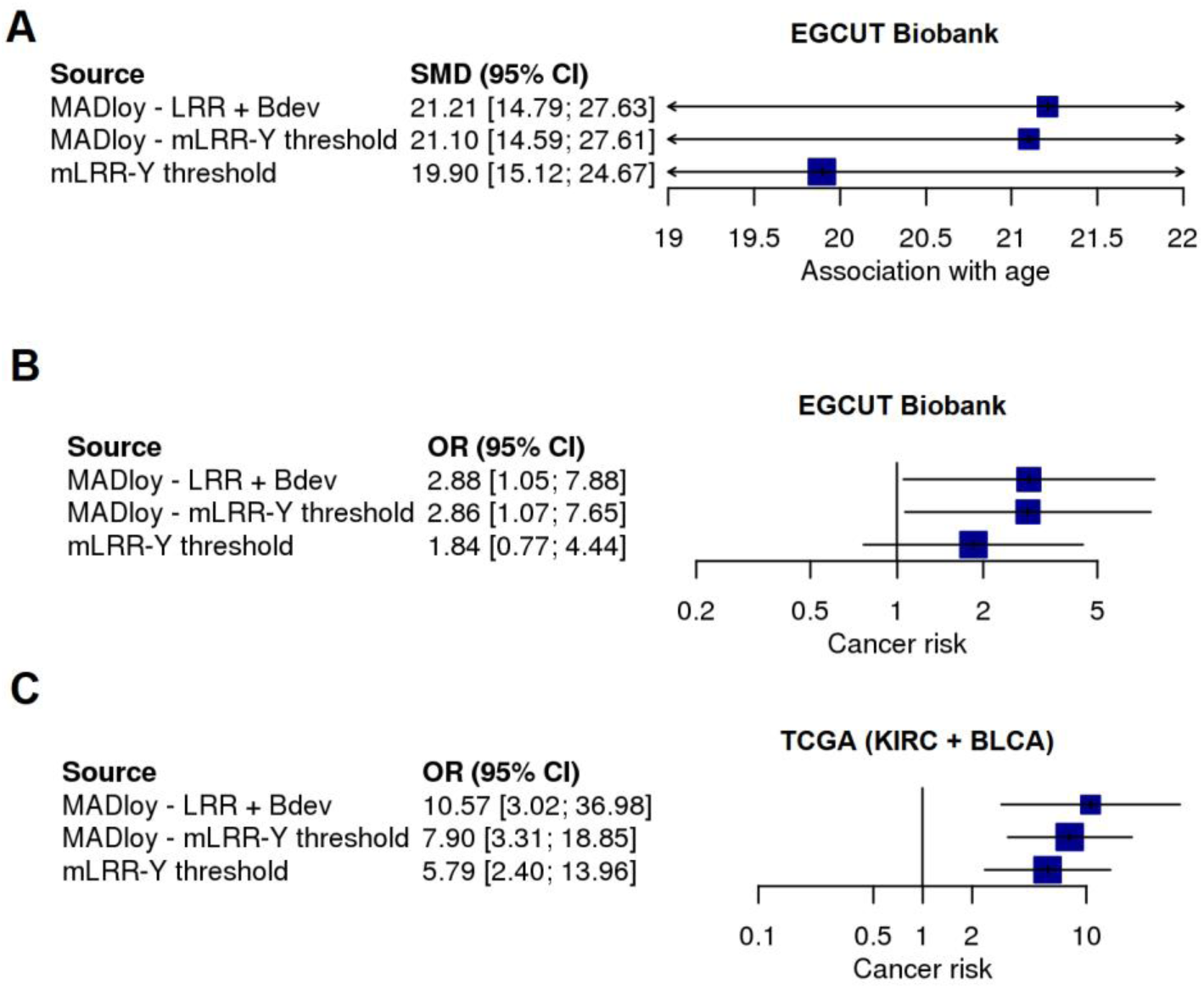
Effect of mLOY misclassification. Effect estimates of three different association studies of continuous (age) and binary phenotypes (cancer) using three different calling methods. Two of them use only LRR information and two different threshold methods and one incorporates the information about B-deviation (Bdev). We observe as the effect is far from the null when using MADloy with both LRR and Bdev.

We also assessed the different mLOY calling procedures in two cancer studies. We studied the differences in frequency of mLOY status between cancer and constitutional tissues of 346 samples from The Cancer Genome Atlas (TCGA) project. The samples belong to Clear Cell Carcinoma (KIRC) and Bladder Urothelial Carcinoma (BLCA) studies. The genotype data comprised 121 constitutional and 103 tumor samples. We first observed that *MADloy* detected *29* samples with mLOY (8,38%) while 34 were detected by *mLRR-Y*_*thresh*_. Thus, *mLRR-Y*_*thresh*_ shows a more liberal approach to mLOY detection, even under presence of 2 GOY samples (0.6%) that are considered as part of the reference population. However, the GOY samples were called by *MADloy* showing the expected increase in mLRR-Y together with a clear BAF split in the PAR1 and PAR2 regions of chromosome X (**Figure S4**).

We then tested the association with cancer status. We observed again that strongest association was when mLOY was called with the full *MADloy* method (OR=10.6, P=4.65×10^−6^). When mLOY was called with *mLRR-Y*_*thresh*_, associations were also significant but their P-values were higher in one order of magnitude (P=1.48×10^−5^ and P=1.59×10^−5^, respectively), in line with previous results and giving further evidence that a more efficient use of the SNP intensity signals in mLOY calling can increment the power to detect associations with phenotypes (**Figure 3B)**. We also tested the association of tumor status with mLOY as a continuous variable (beta=−2.32, P=9.2×10^−5^), and with the cellularity of mLOY obtained by mLRR-Y_cellularity_ (14) (beta=0.98 P=1.59×10^−5^). We therefore observed that while all results were consistent, the new method implemented in MADloy was the most powerful (e.g. showed the most significant p-value).

### Application of MADloy on finding transcriptomic signatures of mLOY

Differences in statistical power become crucial when studying the mechanisms underlying mLOY using genomic and transcriptomic data, as few misclassifications can lead to not detecting a relevant genome or transcriptome-wide association. We therefore apply *MADloy* to comparatively analyzed transcriptomic data in cells from individuals with and without mLOY in blood in the EGCUT study and in cancer tissues in the TCGA study.

We searched for transcription correlates with mLOY in blood using the EGCUT study. We analyzed microarray expression data on 682 individuals (compared with age matched controls with no mLOY) and observed 49 deregulated genes p-value <10^−3^, five of which were significant with p-value < 10^−5^. These included *MED16, HNRNPKP1, TXNDC17, TMEM154*, and *CSF2RA*, the only gene in chromosome Y (PAR1). An enrichment analysis using KEGG and GO databases showed several KEGG pathways that were significantly enriched in differentially expressed genes (**Table 2**). Meaningful pathways including DNA replication (p-value=0.004), mismatch repair (p-value=0.02) and homologous recombination (p-value=0.02) were found significant. Interestingly, only two genes located in the Y chromosome (*CSF2RA* and *EIF1AY)* were among the significantly deregulated gene in blood.

**Table 2.**
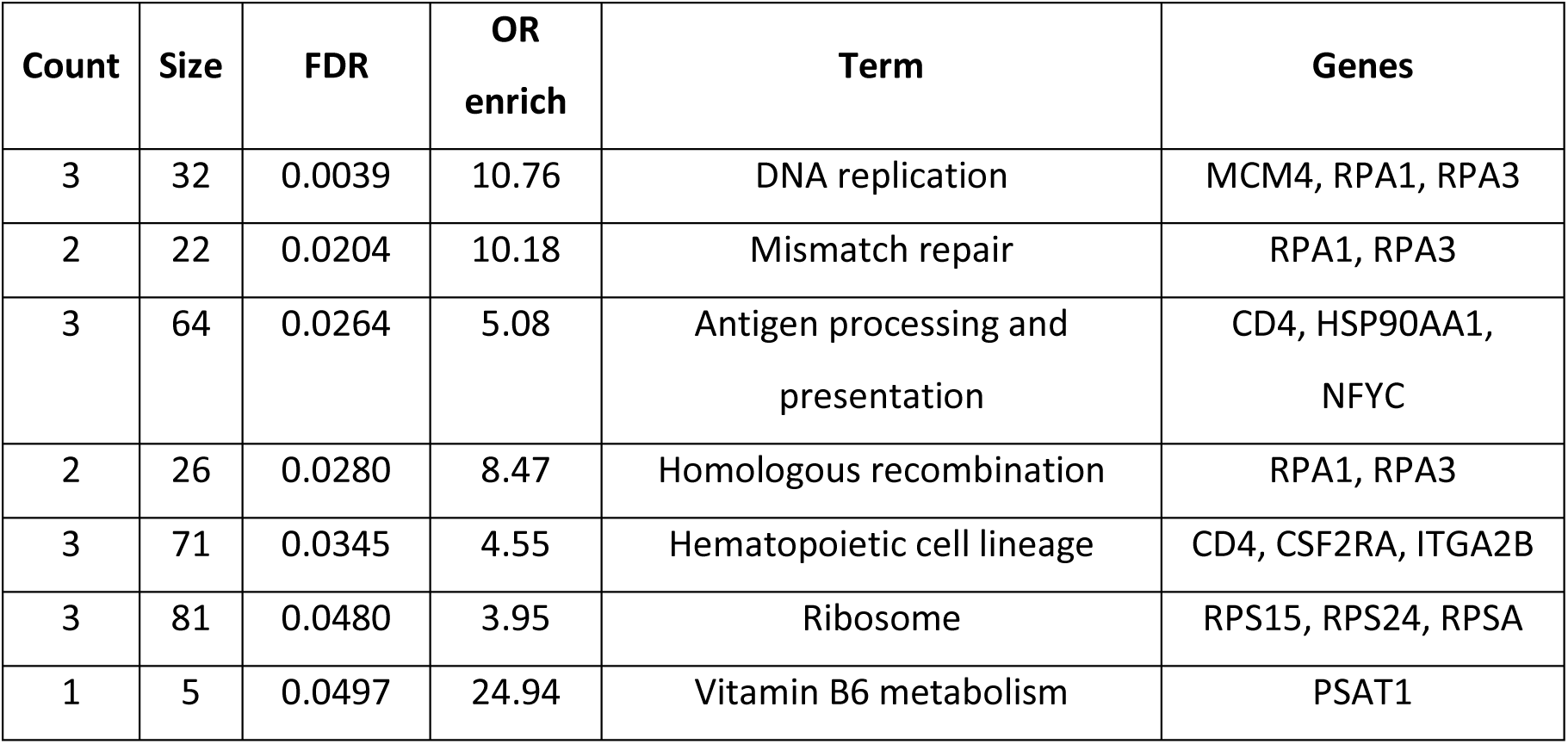
Enrichment analysis of blood transcriptomic data in EGCUT samples. Table includes KEGG categories that are over-enriched using differentially expressed genes at false discovery rate (FDR) lower than 0.05. Only terms with size>=5 are shown.

| Count | Size | FDR | OR enrich | Term | Genes |
| --- | --- | --- | --- | --- | --- |
| 3 | 32 | 0.0039 | 10.76 | DNA replication | MCM4, RPA1, RPA3 |
| 2 | 22 | 0.0204 | 10.18 | Mismatch repair | RPA1, RPA3 |
| 3 | 64 | 0.0264 | 5.08 | Antigen processing and presentation | CD4, HSP90AA1, NFYC |
| 2 | 26 | 0.0280 | 8.47 | Homologous recombination | RPA1, RPA3 |
| 3 | 71 | 0.0345 | 4.55 | Hematopoietic cell lineage | CD4, CSF2RA, ITGA2B |
| 3 | 81 | 0.0480 | 3.95 | Ribosome | RPS15, RPS24, RPSA |
| 1 | 5 | 0.0497 | 24.94 | Vitamin B6 metabolism | PSAT1 |

Transcriptome-wide association analysis of mLOY in TCGA was also performed with RNA-seq data in two different tumor datasets, the KIRC (Kidney renal clear cell carcinoma) and BLCA (Bladder urothelial carcinoma) studies. In both tumors, we found that 8 of the top-10 down-regulated genes by mLOY where in chromosome Y: *TTTY4C, UTY, TMSB4Y, USP9Y, ZFY, EIF1AY, RSP4Y1* and *TTTY15* (adjusted p-values lower than 1.3×10^−28^) (**Figure 4**). This strong correlation with expression is fully supportive of the consistency of mLOY calling by *MADloy*, and indicates that mLOY in tumors leads to a detectable drop in transcription or extreme down-regulation of several genes across the chromosome Y.

**Figure 4.**
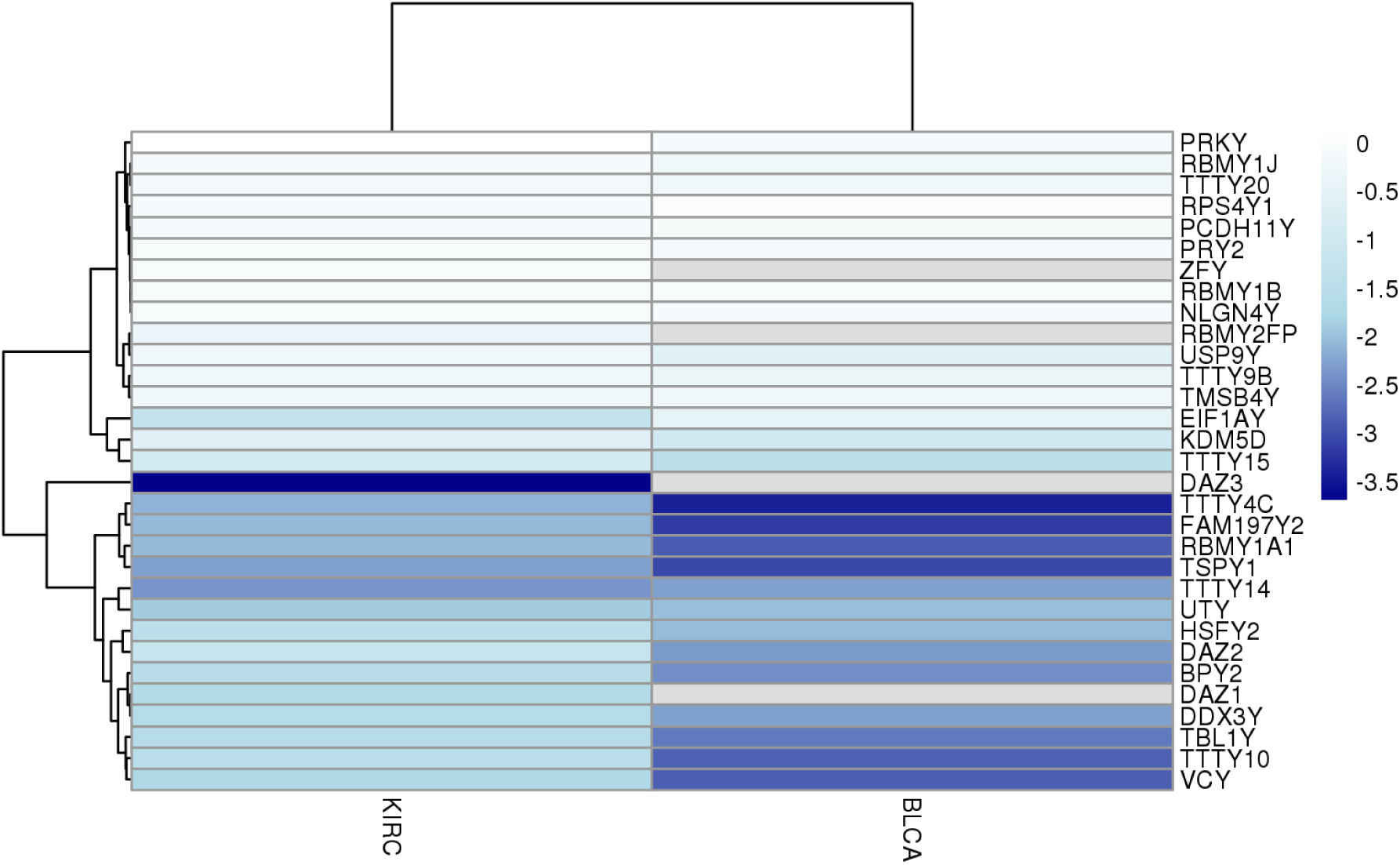
Transcriptomic effects in tumor samples (TCGA). Downregulation of genes in chromosome Y when comparing samples with and without mLOY in KIRC and BLCA datasets of TCGA project.

## Discussion

mLOY is emerging as an important marker of disease for aging men. Forthcoming studies will reveal the extent to which this biomarker can be used to design specific treatment to men. Meanwhile, substantial differences in mLOY calling have arises from the variety of in-house methods used. Here, we have implemented a software in Bioconductor’s framework that incorporate all the current methods described in the literature together with a new robust approach. We show that significant improvements in the detection accuracy of mLOY status from SNP array intensity data can be obtained by using a common matrix-tissue and optimization of the calling methods, by improving the calling threshold in mLRR-Y signal and accounting individuals with chromosome Y gains. Previous methods assumed symmetry in the mLRR-Y distribution and included individuals with gains in the reference population, affecting the threshold to detect mLOY status (1). We observed that individuals with GOY are detectable, in particular in tumor tissues. Therefore, not accounting for asymmetry in the distribution can increase mLOY false positives, reducing the statistical power in the associations due to misclassification. Interestingly we also observed that treating mLRR-Y as a continuous variable did not substantially improved the associations (11). This is likely due to its multimodal nature of its distribution where individuals fall sharply into the normal or mLOY categories.

An additional gain in detection accuracy was observed by an independent analysis of the B-deviation signal (19). While improvements were modest for the associations with age and cancer risk for mLOY in blood, we observed substantial increments in cancer tissues. The inclusion of this B-deviation as an additional signal in mLOY calling allowed us to detect instances where samples are likely contaminated and to confirm individuals with GOY. We have shown that samples classified as mLOY using mLRR-Y with discordant B-deviation values should be visually inspected to define contamination or other non-LOY events. Removing these cases from the analyses in a quality control phase will reduce misclassification errors and improve statistical power. In addition, greater accuracy of mLOY detection is needed if mLOY is to be considered as an individual marker of disease risk.

It has been suggested that discrepancies in associations between mLOY and disease risk or mortality could also be related to the different tissue source of the DNA, despite significant correlation of mLOY calling in blood and buccal derived DNA from the same individuals (9). We show here that the cellularity of mLOY in blood tends to increase with time, but the concordance with mLOY in buccal smear is only 41%, with lower cellularity in buccal samples in most cases. Therefore, different tissue source could lead to differences in mortality ratios and other associations to mLOY, as previously reported (9, 11). On the other hand, the presence of mLOY across different tissues does suggest a common origin in each individual, consistent with a genetic predisposition that has also been documented by genome-wide association studies (7, 8).

Interestingly, the transcriptomic associations of mLOY are quite different in tumor samples from non-tumor blood samples. In kidney and bladder tumors, extreme down-regulation of the entire Y is the main finding, a logical consequence of the decreased number of chromosome Y copies. However, in blood, only two Y chromosome genes are among the most deregulated ones, and deregulation of autosomal genes mainly affects DNA replication, mismatch repair and homologous recombination pathways. This finding could be related to an increasingly oligoclonal leukocyte cell population in people with mLOY, which is consistent with the possibly compromised immune cell function of circulating leukocytes in people with mLOY as the risk factor for disease (1).

The mLOY is increasingly recognized as a male specific risk factor for multiple diseases and, therefore, its interest in personalized treatment and risk management is likely to increase in the coming years. Specific disease and mechanistic studies require a robust estimation of the mLOY status of individuals. SNP intensity data is already available in large epidemiological studies where the effects of environmental conditions can also be investigated. The ability to call mLOY reliably using these data will increase the reproducibility of the findings. We show that *MADloy* is able to replicate the associations of mLOY in blood with cancer and aging while increasing statistical power obtained by current approaches. The method and the bioinformatics tool presented in this work are easy to use and scalable to large studies. Therefore, with *MADloy* we aim to facilitate comparable studies with large number of individuals to better define the role of mLOY in complex diseases and its underlying mechanisms. *MADloy* allows the re-analysis of thousands of existing GWAS data in public available repositories.

## Funding

This research has received funding from the Spanish Ministry of Economy and Competiveness (RTI2018-100789-B-I00). LAPJ lab is funded by the Catalan Department of Economy and Knowledge (SGR2014/1468, SGR2017/1974 and ICREA Acadèmia), and also acknowledges support from the Spanish Ministry of Economy and Competiveness “Programa de Excelencia María de Maeztu” (MDM-2014-0370).

## Conflict of Interest

The authors declare no conflict of interest.

